# Growth deficiency and enhanced basal Immunity in *Arabidopsis thaliana* mutants of *EDM2*, *EDM3* and *IBM2* are genetically interlinked

**DOI:** 10.1101/2023.09.06.556496

**Authors:** Jianqiang Wang, Thomas Eulgem

## Abstract

Mutants of the *Arabidopsis thaliana* genes, *EDM2*, *EDM3* and *IBM2* are known to show defects in a diverse set of defense and developmental processes. For example, they jointly exhibit enhanced levels of basal defense and stunted growth. Here we show that these two phenotypes are functionally connected by their dependency on the salicylic acid biosynthesis gene *SID2* and the basal defense regulatory gene *PAD4*. Stunted growth of *edm2*, *edm3* and *ibm2* plants is a consequence of up-regulated basal defense. Constitutively enhanced activity of reactive oxygen species-generating peroxidases, we observed in these mutants, appears also to contribute to both, their enhanced basal defense and their growth retardation phenotypes. Furthermore, we found the histone H3 demethylase gene *IBM1*, a direct regulatory target of EDM2, EDM3 and IBM2, to be at least partially required for the basal defense and growth-related effects observed in these mutants. We recently reported that *EDM2*, *EDM3* and *IBM2* coordinate the extent of basal immunity with the timing of the floral transition. Together with these observations, data presented here show that at least some of the diverse phenotypic effects in *edm2*, *edm3* and *ibm2* mutants are genetically interlinked and functionally connected.

## Introduction

Sustained or constitutive activation of immune responses in plants is often associated with reduced growth and other developmental abnormalities, a phenomenon related to the concept of defense-growth trade-off. It has long been believed that such effects are due to competition of immunity-related and developmental processes for limited metabolic resources [1]. When plants are attacked by microbial pathogens, metabolic resources are preferentially allocated to defense, resulting in enhanced immunity, but at the expense of reduced growth [2,3].

The *Arabidopsis thaliana* (Arabidopsis) Plant Homeodomain (PHD) finger protein-encoding *EDM2* gene and the RNA Recognition Motif (RRM) domain protein-encoding genes *EDM3/AIPP1* (hereafter *EDM3*) and *IBM2/ASI1/SG1* (hereafter *IBM2*) are know to jointly affect multiple immunity-related and developmental processes [4–12]. They cooperate in a strong race-specific pathogen defense mechanism mediated by the disease resistance gene *RPP7*, that encodes an NLR-type immune receptor [13–15]. They also work together in an unspecific immune response effective against a wide range of different pathogens, termed basal immunity or basal resistance [16,17]. While they promote *RPP7*-mediated immunity, *EDM2*, *EDM3* and *IBM2* suppress basal defense. The latter process is coordinated by these three genes with the floral transition [17]. Besides this and other developmental roles, *EDM2*, *EDM3* and *IBM2* promote overall growth, as their mutants exhibit reduced fresh weight and smaller rosette leaves [5,11].

Mircoarray and RNA-seq studies with their mutants have shown *EDM2*, *EDM3* and *IBM2* to affect transcript levels of large overlapping sets of genes and transposable elements, while epigenome profiling revealed joint effects on cytosine methylation and/or the repressive histone mark H3K9me2 at numerous chromatin sites [5, 8–10,12,16]. By chromatin-immunoprecipitation (ChIP) the EDM2, EDM3 and IBM2 proteins were found to co-localize at some of these loci [7–10,12,13,16,18]. *RPP7* and the histone H3K9 demethylase gene *IBM1* are among the direct target sites of EDM2, EDM3 and IBM2. Upon co-recruitment to heterochromatic stretches at each of these two (and several other) target loci, they prevent premature transcript termination at alternative polyadenylation signals, thereby promoting the synthesis of full-length mRNAs [8–10,12,13,15,16]. Consistent with their co-localization at chromatin sites, physical interactions among these three proteins were observed. While both EDM2 and IBM2 appear to directly interact with EDM3, interactions between EDM2 and IBM2 seem indirect and require EDM3 as a molecular bridge [12]. Recent results strongly suggested that EDM3/IBM2 interactions can be isoform specific [17]. Due to alternative splicing, two distinct protein isoforms are expressed for each of these two genes; a shorter one (EDM3S; IBM2S) and a longer one (EDM3L; IBM2L). Regarding their role in the suppression of basal defense and the promotion of the floral transition, only the two longer isoforms, EDM3L and IBM2L, and not the shorter EDM3S and IBM2S isoforms, cooperate [17].

Consistent with their role in suppressing basal defense, we observed that large overlapping sets of defense-related genes and immune receptor genes are constitutively transcriptionally up-regulated in mutants of *EDM*2, *EDM3* and *IBM2* [15–17]. Thus, *edm2*, *edm3* and *ibm2* mutants exhibit three typical effects common to many other Arabidopsis mutants with constitutively elevated levels of basal immunity [19–23]: (1) constitutively elevated expression of defense genes, (2) enhanced basal defense and (3) retarded growth. Here we show that enhanced basal defense and growth retardation in *edm2*, *edm3* and *ibm2* mutants are genetically coupled by their dependency on the salicylic acid-associated defense genes *SID2* and *PAD4*, implying that the latter effect is likely a consequence of the former one. We further present data suggesting that increased peroxidase activity, as a part of the constitutive basal defense response, is a partial cause of reduced *edm2*, *edm3* or *ibm2* growth rates. In addition we found that the direct EDM2, EDM3 and IBM2 target gene *IBM1*, is partially required for the basal defense and growth-related phenotypes. Each of these effects is mediated by the longer isoforms of EDM3 and IBM2 but not the shorter ones. Together with our previous report on the coordination of basal defense with the timing of the floral transition by *EDM2*, *EDM3* and *IBM2*, our new results show that these three chromatin-associated regulators link various defense and developmental processes, likely contributing to a balanced defense-growth trade-off that enables efficient developmental transitions and limits growth penalties due to energetically costly basal immune responses.

## Results

### *EDM2*, *EDM3L* and *IBM2L* positively affect plant growth

Previously, we showed that *edm2* mutants exhibit decreased fresh weight and rosette leaf expansion compared to their wild type parental lines [5]. Whole plant fresh weight and rosette leaf expansion are also decreased in mutants of *EDM3* and *IBM2* (Fig 1A, 1B and S1 Fig). We used the previously described *sg1-3* mutant allele of *IBM2* [11], *edm2-2* allele of *EDM2* [13] and *edm3-2* of EDM3 [17] for all experiments in this study. We further found that *edm2*, *edm3*, and *ibm2* mutants have shorter primary roots (Fig 1C and S1 Fig). Moreover, only the longer isoforms of EDM3 and IBM2 we previously described, EDM3L and IBM2L [17], can rescue each of these two growth-related defects in *edm3* or *ibm2* mutants, respectively. Functional complementation by the shorter isoforms EDM3S and IBM2S failed as the respective transgenic lines exhibit similar phenotypes as their parental mutants (S1 Fig). Overall, these data show that EDM3L and IBM2L, like EDM2, positively affect overall plant growth.

**Fig 1.**
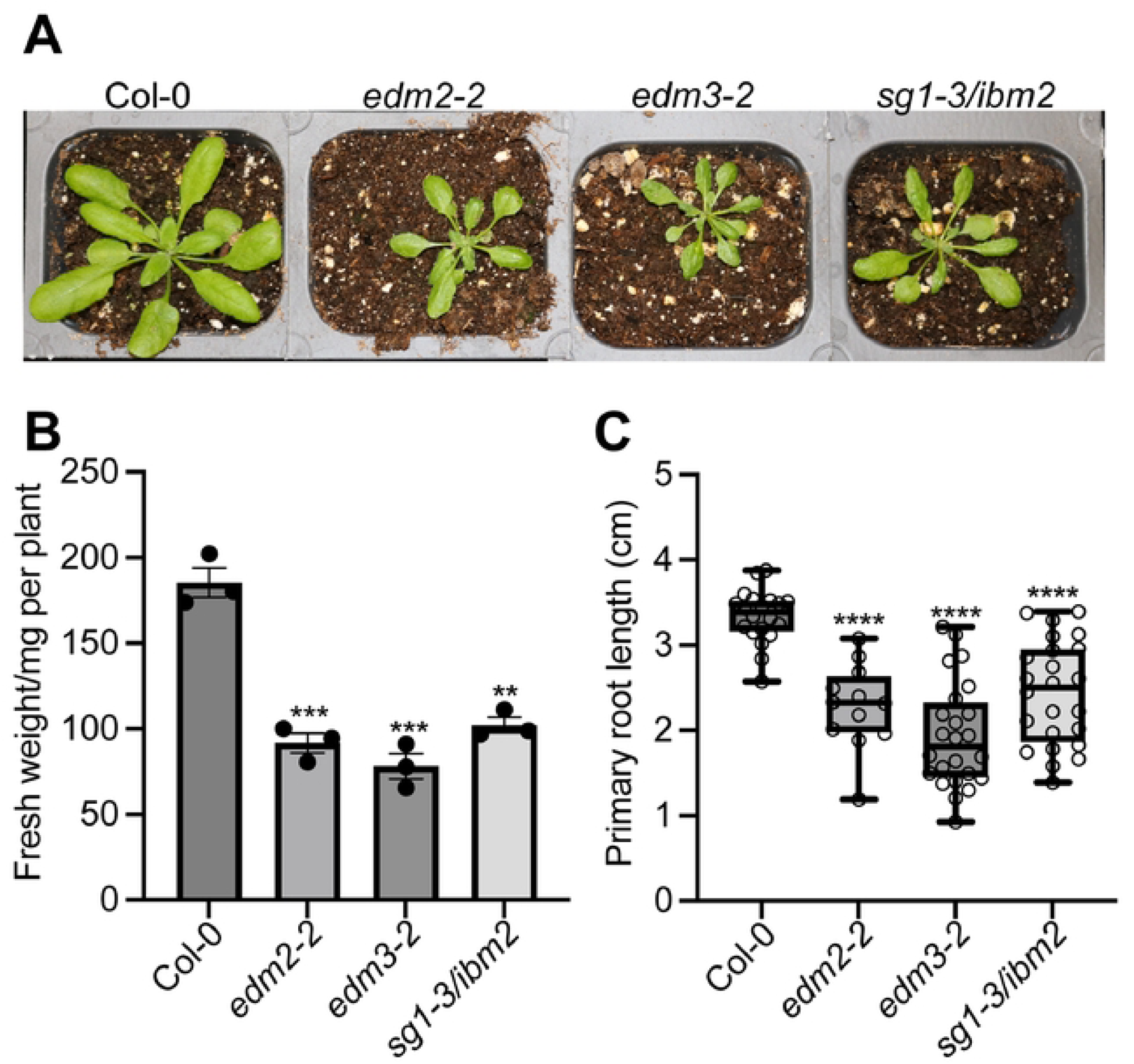
*EDM2*, *EDM3* and *IBM2* positively affect overall plant growth. **A.** Representative images of 25-day-old Arabidopsis plants of the indicated genotypes grown in soil. **B.** Average fresh weight of 25-day-old whole plants of the indicated genotypes. Error bars represent standard errors from three independent experiments, each with eighteen plants. **C.** Primary root length of 7-day-old plants of the indicated genotypes grown on agar plates. Primary root lengths of more than twelve plants of each genotype were measured using ImageJ. Data information: Asterisks indicate significant differences compared to Col-0 based on Student’s t-test. (**, *p* < 0.01; ***, *p* < 0.001; ****, *p* < 0.0001).

### Enhanced Basal Immunity in *edm2*, *edm3* and *ibm2* suppresses plant growth

To examine if growth defects in *edm2*, *edm3* and *ibm2* mutants are caused by their constitutively enhanced basal immunity, we used Arabidopsis lines with defects in defense responses controlled by salicylic acid (SA). The *sid2-2* mutant is compromised in the defense-associated biosynthesis of this phytohormone [24], while the transgenic *NahG* line cannot accumulate SA, due to the expression of a bacterial SA hydroxylase gene [25]. The *pad4-1* mutant is deficient in a SA-responsive signaling step [26]. We crossed *edm3* and *ibm2* mutants to each of these three lines and performed experiments with lines homozygous for each altered component (mutation or transgene). In addition, we generated *edm2-2;sid2-2* and *edm2-2;pad4-1* double mutants. We first examined if enhanced immunity in *edm2*, *edm3* and *ibm2* is reduced by blockage of SA-mediated defense. We tested Arabidopsis seedlings one week after infection with the Noco2 isolate of the pathogenic oomycete *Hyaloperonospora arabidopsidis* (*Hpa*) infection, which is virulent on wild type Col-0 plants and triggers basal defense in this accession [27]. As shown previously [17], *edm2*, *edm3* and *ibm2* plants show reduced susceptibility against *Hpa*Noco2 compared to Col-0, indicating that basal defense in these single mutants is enhanced (Fig 2A-D). However, we observed complete loss of this enhanced basal defense phenotype in all double mutants as well as the *edm3-2;NahG* and *sg1-3/ibm2;NahG* lines.

**Fig 2.**
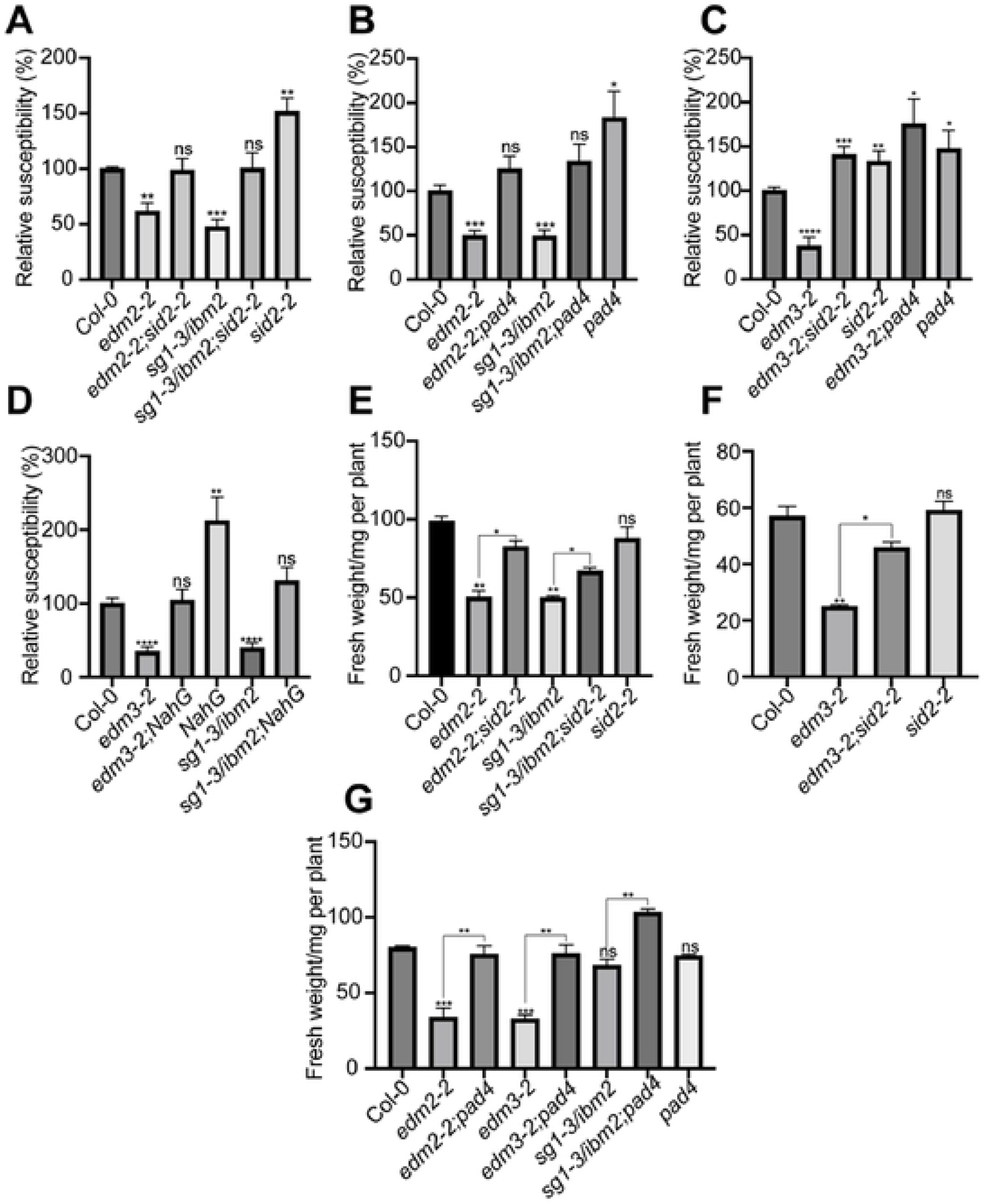
Enhanced basal immunity in mutants of *EDM2*, *EDM3* and *IBM2* requires salicylic acid, *SID2,* and *PAD4*. **A-D.** Enhanced basal immunity in *edm2*, *edm3* and *ibm2* mutants was disrupted by introducing the *sid2-2* or *pad4* mutations or the *NahG* transgene. Two-week-old seedlings were sprayed-inoculated with 3 x 10^4^/ml *Hpa*Noco2 spores. *Hpa*Noco2 spores were counted one week post infection, which were divided by the corresponding fresh weight and shown as percentage relative to wild type. **E-G.** Fresh weight of 24-day-old whole plants **(E)**, 23-day-old whole plants **(F)** or 25-day-old whole plants **(G)** of the indicated genotypes, which were grown on soil. Data information: Data shown in each separate panel were generated simultaneously. Error bars represent standard errors from three independent experiments. Asterisks indicate significant differences compared to Col-0 based on Student’s t-test (A-D) or by one-way ANOVA with Brown-Forsythe and Welch’s test (E) or by ordinary one-way ANOVA (F & G). (*, p < 0.05; **, p < 0.01; ***, p < 0.001; ****, p < 0.0001; ns, no significance).

We further observed the growth-related effects on whole plant fresh weight and primary root length of *edm2*, *edm3* and *ibm2* mutants to be partly or fully reversed to wild type levels in the respective double mutants with *sid2-2,* or *pad4* (Fig 2E-G and S2 Fig). We made the same observation with *edm3*;*NahG* and *ibm2*;NahG lines (Fig 2E-G; and S2 Fig). One deviation are the fresh weight measurements of *ibm2* plants in Figure 2G, which seem lower than Col-0, but are not significantly different from this wild type reference line. The Student’s t-test *p* value for this is slightly above the significance cutoff. However, compared to Col-0 we observed fresh weight and overall growth of *ibm2* plants to be significantly reduced with very high consistency in all other experiments of this study (Figs 1; 2E; 6B; S1B, E & F; S2A, D & E; S3). Still the fresh weight of *ibm2;pad4* double mutants in this set of experiments is significantly higher than that of the *ibm2* single mutant (Fig 2G).

Overall, these findings show that enhanced immunity in mutants of *EDM2*, *EDM3* and *IBM2* relies on the SA-dependent basal defense pathway. As this pathway is further required for the reduction of whole plant fresh weight and primary root length in *edm2*, *edm3* and *ibm2* plants, we conclude that the growth-related effects are a consequence of the enhanced basal defense phenotype of these mutants.

### *EDM2*, *EDM3* and *IBM2* suppress ROS-generating peroxidase activity by down-regulation of type III peroxidase genes

A reactive oxygen species (ROS)-generating oxidative burst is an early event among plant immune responses [28][19–23][29,30]. In Arabidopsis, NADPH oxidases and cell wall-associated type III peroxidases are two main sources of ROS in the apoplast [31,32]. ROS are not only involved in immune response but also affect plant growth. ROS accumulation can stiffen cell walls by cross-linking its components, thereby inhibiting cell expansion [33]. Due to these dual effects, ROS have been linked to the defense-growth trade-off as a shared component of both processes [34].

Re-inspecting our previous RNA-seq analyses on *EDM2*-mediated transcriptome changes [16] we found 27 type III peroxidase genes and two NADPH oxidase genes, *RbohD* and *RbohF*, to be significantly up-regulated in *edm2* plants compared to their wild type parental line, while the gene encoding the H_2_O_2_ scavenging enzyme, *CAT1*, is significantly down-regulated in this mutant. Since over-accumulation of the ROS H_2_O_2_ can reduce cell wall extensibility [33], the joint upregulation of type III peroxidase and NADPH-oxidase genes may contribute to growth-related effects in *edm2* plants.

We selected 11 of these 27 type-III peroxidase genes, which show either particularly strongly increased transcript levels in *edm2* plants or are possible direct target genes of EDM2 and IBM2 based on previous ChIP-seq analyses [18]. For nine of these 11 genes (except *PRX31* and *PRX62*) we consistently observed in qRT-PCRs constitutively elevated transcript levels in the tested *edm2-2*, *edm3-2* and *sg1-3/ibm2* mutants compared to Col-0 (Fig 3A). While we could not confirm the up-regulation of *RbohD* and *RbohF* (Fig 3B), we observed by qRT-PCR a significant up-regulation of *RbohB,* another member of this family of NADPH oxidase genes in all three tested mutants (Fig 3B).

**Fig 3.**
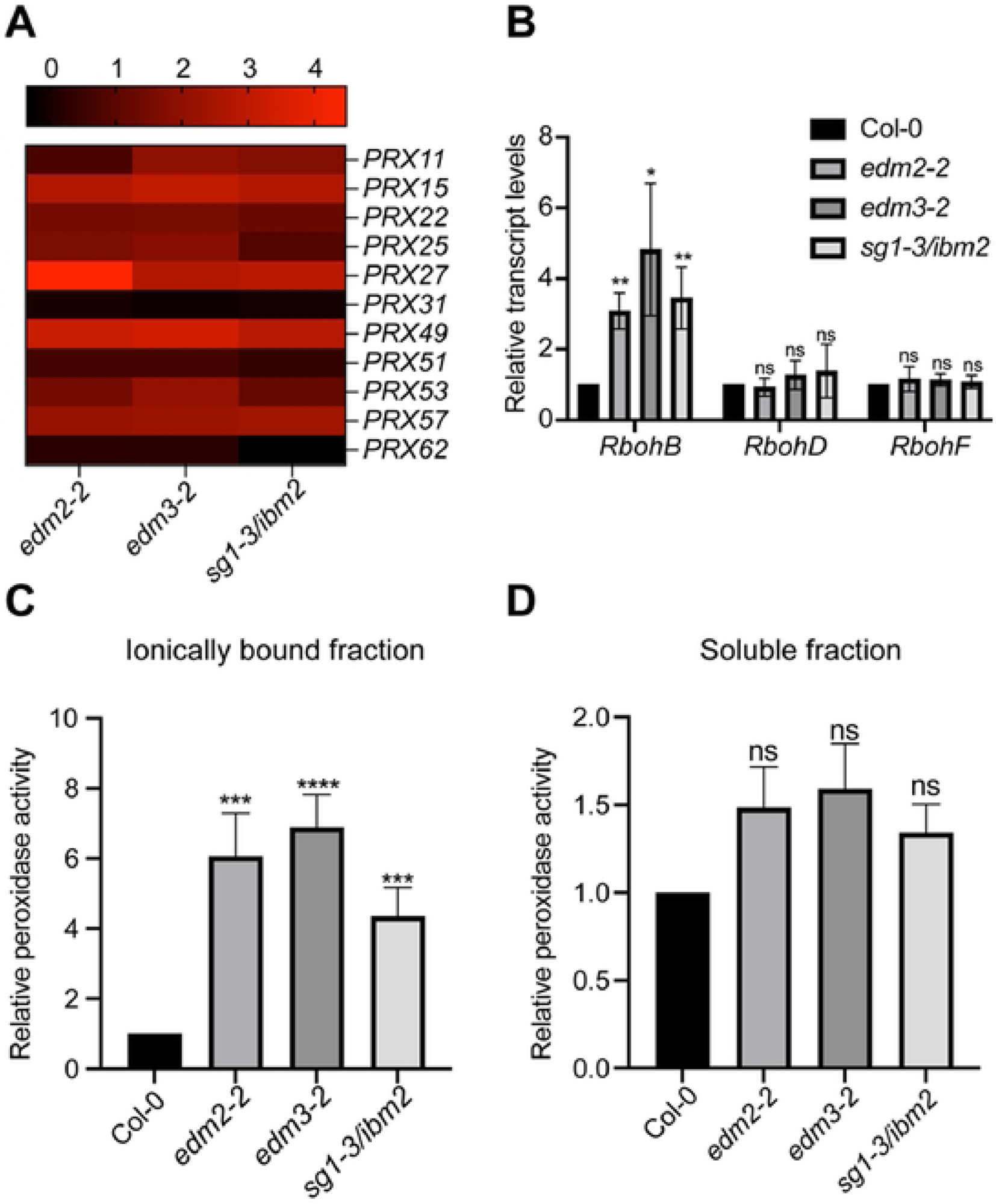
*EDM2*, *EDM3* and *IBM2* suppress expression levels of genes involved in the generation of ROS as well as activity levels of peroxidases. **A.** The transcript levels of selected peroxidase genes in mutants of *EDM2*, *EDM3* and *IBM2* in the aerial parts measured by qRT-PCR. Log2-Fold Change (FD) relative to transcript levels in wild type was used to create this heatmap. The brightest red of the above scale represents the highest increase relative to wild type plants. Numbers above the scale bar represent log2 fold change values of transcript level increases relative to Col-0. **B.** Transcript levels of NADPH oxidase genes in mutants of *EDM2*, *EDM3* and *IBM2* in the aerial parts measured by qRT-PCR. **C & D**. Relative peroxidase activity in mutants of *EDM2*, *EDM3* and *IBM2*. Peroxidase activity was measured in the aerial parts of two-week-old plants, divided by fresh weight and shown relative to the wild type plants. Data information: Error bars represent standard errors from three independent experiments. Asterisks indicate significant differences analyzed by student’s t-test. (*, p < 0.05; **, p < 0.01; ****, p < 0.0001; ns, no significance).

We further tested if peroxidase enzyme activity is also increased in these mutants. Type III peroxidases are known to be localized to the cell wall. Thus, the transcriptional upregulation of these enzymes we observed in mutants of *EDM2*, *EDM3* and *IBM2* should lead to increased peroxidase activity in their ionically bound protein fractions. Indeed, we found that compared to Col-0, the *edm2*, *edm3* and *ibm2* mutants jointly exhibit a noticeable increase in constitutive peroxidase activity in the ionically bound fraction (Fig 3C), but not the soluble fraction (Fig 3D).

To test if the increased peroxidase activity contributes to the growth defects observed in *edm2*, *edm3* and *ibm2* plants, we treated them with 20μM of the peroxidase inhibitor salicylhydroxamic acid (SHAM), at day 14 and day 16 and compared their rosette areas at day 25. We observed a significant increase in rosette areas in all tested mutants after SHAM treatment compared to untreated plants, while there is no difference in wild type after treatment with this inhibitor (Fig 4). Overall, these data suggest that increased peroxidase activity contributes to growth defects in *edm2*, *edm3* and *ibm2* mutants.

**Fig 4.**
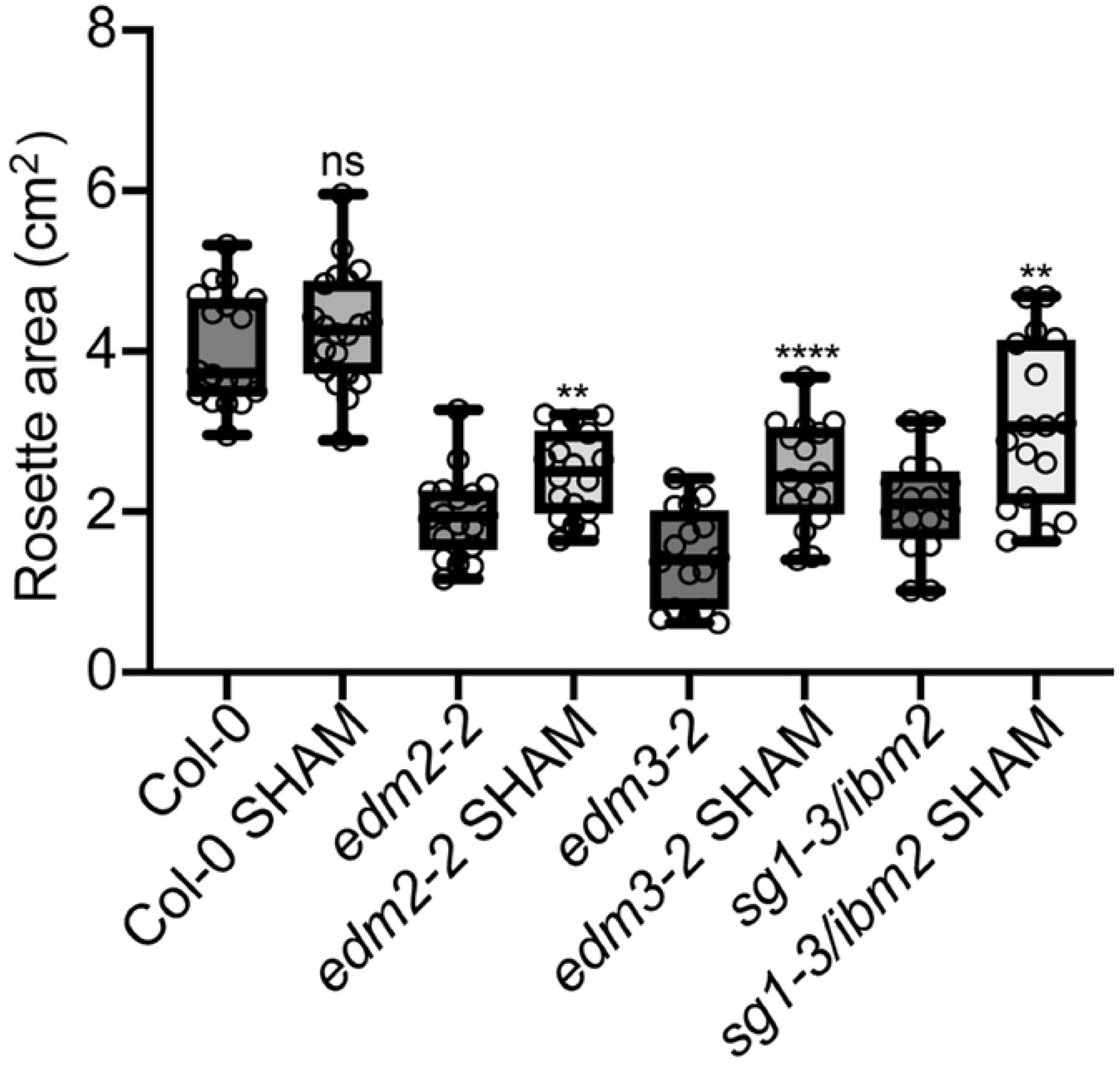
Growth defects in mutants of EDM2, EDM3 and IBM2 are partly rescued by inhibiting peroxidase activity. Rosette areas of 25-day-old plants of the indicated genotypes. 20μM SHAM was applied to soil grown plants at day 14 and day 16. Rosette area (n ≥ 16) was measured using ImageJ. Data information: Asterisks indicate significant differences between SHAM and non-SHAM treated plants of the same genotype based on Student’s t-tests. (*, p < 0.05; **, p < 0.01; ****, p < 0.0001; ns, no significance).

### Basal *Hpa*Noco2 resistance in mutants of *EDM2*, *EDM3* and *IBM2* is decreased by inhibiting peroxidase activity

To examine if enhanced basal resistance in *edm2*, *edm3* and *ibm2* mutants is partially due to the increased type III peroxidase activity, we pre-treated all three mutants with SHAM 24 hours before infection with *Hpa*Noco2. Compared to untreated plants we observed a larger number of *Hp*aNoco2 spores in all three mutant plants after treatment with SHAM and a mild increase in Col-0 wild type plants with SHAM treatment (Fig 5), suggesting increased peroxidase activity contributes to enhanced immunity in all three mutants.

**Fig 5.**
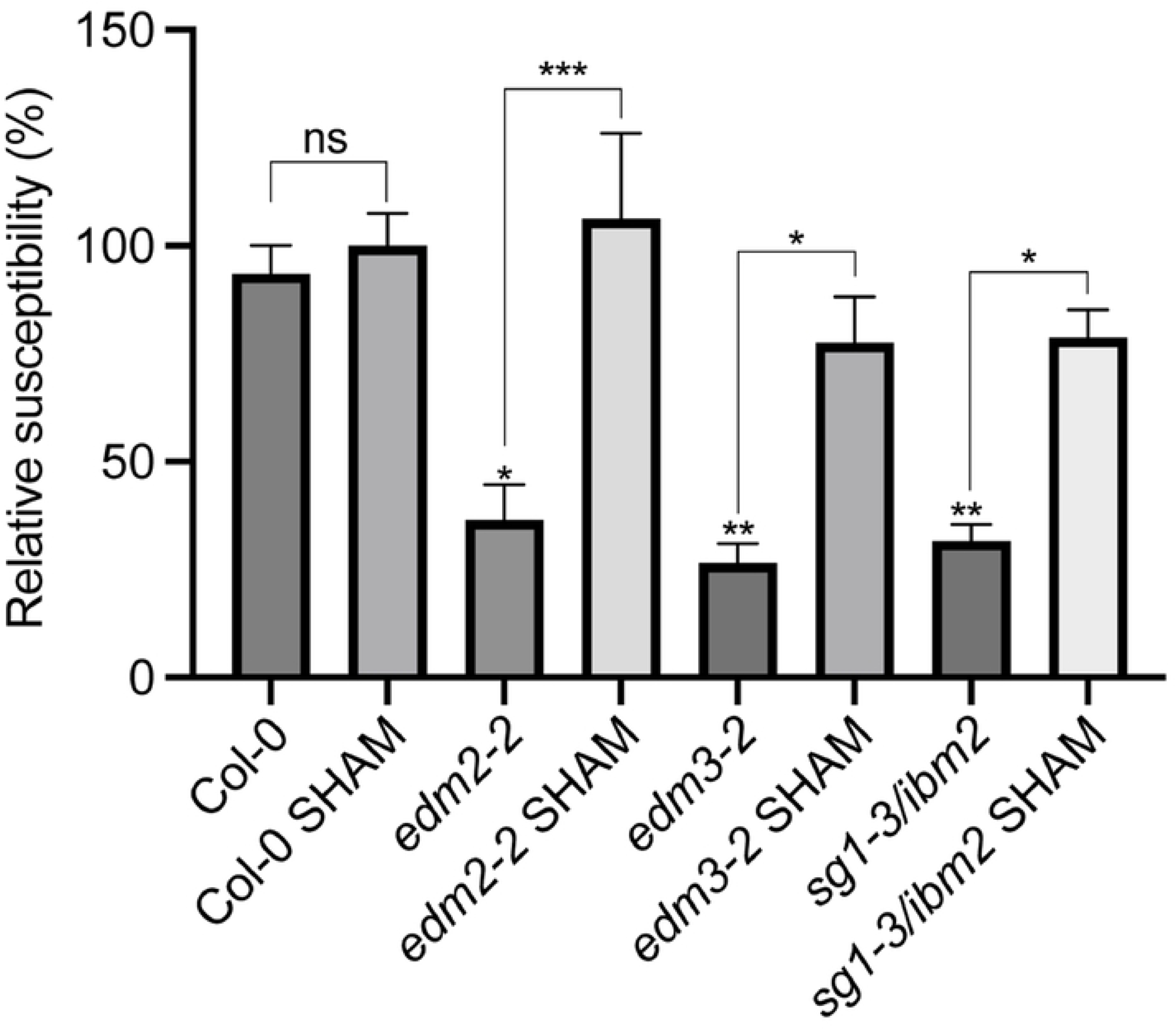
Increased type III peroxidase activity is likely to contribute to enhanced basal defense in mutants of *EDM2*, *EDM3* and *IBM2*. Two-week-old plants of the indicated genotypes were pre-treated with 20μM SHAM 24 hours before sprayed-inoculated with 1 x 10^4^/ml *Hpa*Noco2 spores. *Hpa*Noco2 spores numbers were determined one week after infection. Spore numbers were normalized to the corresponding fresh weights. Data information: Error bars represent standard errors from three independent experiments. Asterisks indicate significant differences between plants of the same genotype based one-way ANOVA. (*, p < 0.05; **, p < 0.01; ***, p < 0.001; ns, no significance).

### IBM1L partly rescues growth defects and disease susceptibility in mutants of *EDM2*, *EDM3* and *IBM2*

At the epigenome and transcriptome levels mutations in the histone H3 demethylase gene *IBM1* phenocopy many effects of *edm2*, *edm3* and *ibm2* mutations. Mutants of *IBM1* are also growth-retarded. *EDM2*, *EDM3* and *IBM2* are known to promote the expression of the functional full-length IBM1 isoform (IBM1L) by suppressing proximal transcript polyadenylation at an alternative polyadenylation signal in a heterochromatic repeat region in the seventh *IBM1* intron [8,9]. To examine dependency of *EDM2*, *EDM3* and *IBM2*-mediated basal defense and growth-related effects on *IBM1*, we transformed into *edm2*, *edm3* and *ibm2* mutants a genomic clone containing the entire *IBM1* gene (*gIBM1*) or a mutated version of this clone with a deletion of the heterochromatic repeats in its seventh intron (*gIBM1ΔHR*). While *gIBM1* was shown before to rescue various phenotypes of *ibm1* mutants, *gIBM1ΔHR*, but not *gIBM1*, rescues low expression levels of the functional full-length IBM1L isoform and leaf abnormalities in *edm2* and *ibm2* mutants [8,9]. The *gIBM1ΔHR* clone lacks the proximal alternative polyadenylation signal in intron 7 and, therefore, does not require *EDM2*, *EDM3* and *IBM2* for proper expression of IBM1L. We observed that *Hpa*Noco2 susceptibility, fresh weight and primary root length were at least partially restored to wild type levels in *edm2*, *edm3* and *ibm2* mutants containing *gIBM1ΔHR*, but not *gIBM1* (Fig 6A & 6B; Fig S3). Compared to Col-0 in most cases *Hpa*Noco2 susceptibility levels, fresh weight and primary root length were still significantly reduced in the mutants containing *gIBM1ΔHR*. However, in all cases these values were also significantly increased compared to their respective *gIBM1*-containing counterparts. Thus, the histone H3 demethylase gene *IBM1*, is at least partially responsible for *EDM2*, *EDM3* and *IBM2*-mediated suppression of basal defense and the resulting promoting effects on growth.

**Fig 6.**
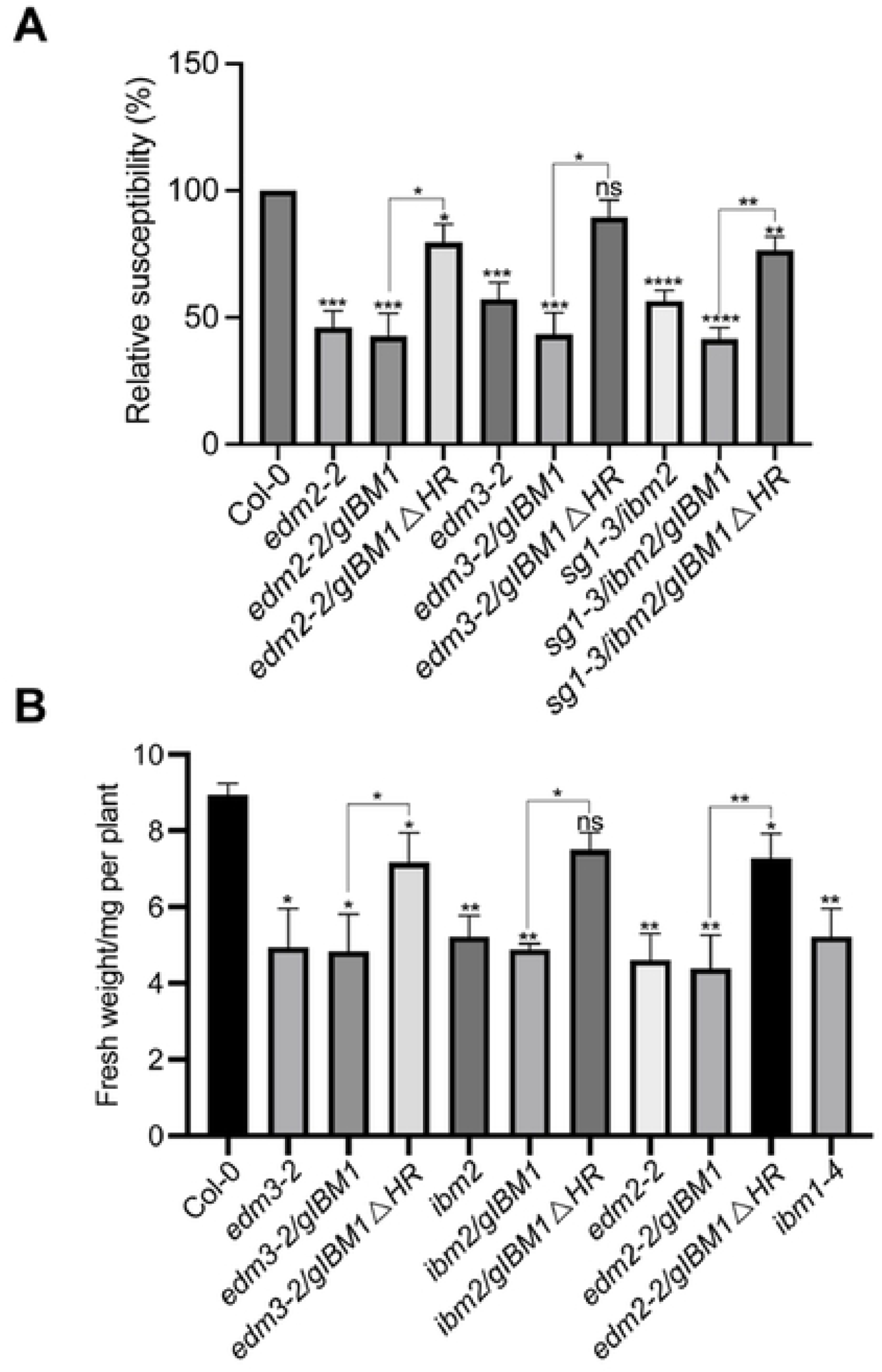
Growth defects and *Hpa*Noco2 susceptibility were partly rescued by IBM1L. **A.** *Hpa*Noco2 susceptibility is rescued in mutants of *EDM2*, *EDM3* and *IBM2* by a genomic *IBM1* clone with a deletion of its heterochromatic repeats in intron 7 (*gIBM1ΔHR*), but not by the complete genomic region lacking the deletion (*gIBM1*). Two-week old plants were spray-inoculated with 3 x 10^4^/ml *Hpa*Noco2 spores. *Hpa*Noco2 spores were counted one week post infection. The resulting spore numbers were divided by the respective fresh weight and shown as percentage relative to wild type. **B.** Fresh weight of twelve-day-old whole plants of the indicated genotypes. Data information: Error bars represent standard errors from three independent experiments. Asterisks indicate significant difference compared to Col-0 based on Student’s t-test (*, p < 0.05; **, p < 0.01; ***, p< 0.001; ****, p < 0.0001, ns, no significance).

## Discussion

Numerous Arabidopsis mutants with constitutively activated immunity have been described before, such as *cpr (<u>c</u>onstitutive expresser of <u>PR</u> genes)* [22], *cim (<u>c</u>onstitutive <u>im</u>munity)* [23], *lsd (lesion-<u>s</u>imulating <u>d</u>isease resistance)* [20], *acd* (*<u>a</u>ccelerated <u>c</u>ell <u>d</u>eath)* [35], *dnd* (*<u>d</u>efense <u>n</u>o <u>d</u>eath)* [36] and *edr* (enhanced disease resistance) [37] mutants. Common to mutants of these classes are constitutively enhanced expression levels of defense-related genes, enhanced basal defense and retarded growth or other developmental defects. These effects are typically dependent on SA-mediated defense signaling. Mutants of *EDM2*, *EDM3* and *IBM2* exhibit the same set of phenotypic effects. Using double mutants we showed enhanced basal defense and retarded rosette and root growth of *edm2*, *edm3* and *ibm2* plants to depend on the SA biosynthesis gene *SID2* and the SA signaling gene *PAD4*. Expression of the bacterial SA hydroxylase gene *NahG* leads to the same effects in *edm3* and *ibm2* plants. Thus, both the enhanced basal defense and retarded growth phenotypes of the *edm2*, *edm3* and *ibm2* mutants are genetically interlinked. As blockage of SA-mediated defense responses rescues the growth defects of these mutants, their growth retardation must be a consequence of the enhanced basal defense. Consequently *EDM2*, *EDM3* and *IBM2* contribute to the defense-growth trade of by limiting the extent of energetically costly basal defense responses, thereby, prioritizing overall growth. We recently showed the coordination of basal defense with the timing of the floral transition to be mediated by the respective longer isoforms of EDM3 and IBM2. This also applies to the growth promoting effects, which is consistent with the view that these processes are interlinked.

While *EDM2*, *EDM3* and *IBM2* suppress basal defense, they have the opposite effect on immunity mediated by the NLR-type immune receptor genes *RPP7* and *RPP4*, the expression of which they promote [4,12,14–17]. In particular *RPP7*-mediated immunity may be energetically costly to plants, as this defense mechanism is unusually strong leading to very tight disease resistance [4,39]. In addition, mutants of *RPP7* exhibit enhanced overall growth, suggesting that even in a resting state this immune receptor consumes energy-related resources (Yan Lai, Tokuji Tsuchiya; Jianqiang. Wang, and Thomas. Eulgem, unpublished). Thus, the contribution to the defense-growth trade-off of *EDM2*, *EDM3* and *IBM2* seems complex. Rather than serving as general suppressors immune responses, *EDM2*, *EDM3* and *IBM2* seem to mediate a certain balance of defense-related processes, by having counter-directional effects and promoting some defense mechanisms at the expense of others. In addition, they coordinate the extent of basal defense with at least one important developmental transition, the one from vegetative growth to flowering.

As a likely joint contributor of the elevated basal immunity in *edm2*, *edm3* and *ibm2* plants we identified constitutively increased activity of type III peroxidases, a class of enzymes known to contribute to the defense-associated oxidative burst. The oxidative burst is a very early immune response, occurring within minutes when plants are invaded by pathogens [28]. Plant defenses require ROS generated in the oxidative burst, which serve as defensive antimicrobial compounds, cross-linkers of cell wall macromolecules as well as defense signaling molecules [40,41]. Expression of a set of type III peroxidase-encoding genes is constitutively upregulated in *edm2*, *edm3* and *ibm2* mutants. Consistent with the known localization of type III peroxidases to plant cell walls [32], we observed increased peroxidase activity in anionic protein fractions and not in soluble protein fractions of *edm2*, *edm3* and *ibm2* plants. Blockage of peroxidase activity by the inhibitor SHAM partially reversed the basal defense and growth levels of these mutants to those observed in their wild type parents. Thus, increased type III-mediated accumulation of ROS may contribute to the enhanced basal defense phenotype observed in *edm2*, *edm3* and *ibm2* plants. Apoplastic ROS accumulation may also reduce cell wall extensibility possibly due to lignin formation and cross-linking of other macromolecular components [33,42], thereby suppressing plant cell growth. However, using standard H_2_O_2_ detection assays, we were unable to detect any significant increase of this reactive oxygen species in *edm2*, *edm3* and *ibm2* plants. Perhaps the actual increase of ROS is only incremental and too weak to be detectable by the assays we applied.

Many epigenomic and transcriptomic effects caused by *EDM2*, *EDM3*, *IBM2* are known to be mediated by their direct target gene *IBM1*. By suppressing proximal polyadenylation of IBM1 transcripts *EDM2*, *EDM3* and *IBM2* jointly promote expression of the functional full-length IBM1L isoform. Here we found suppression of basal defense as well as the promotion of rosette and root growth by *EDM2*, *EDM3* and *IBM2* to depend at least partially on IBM1L. We already reported overlapping effects on the expression of defense-related and immune receptor genes by *EDM2* and *IBM1* [16]. However, we also observed that each of these regulators affects transcript levels of separate sets of defense-related and immune receptor genes. *IBM1*, in contrast to *EDM2*, *EDM3* and *IBM2*, does not affect *RPP7* expression and immunity-mediated by this NLR gene.

Collectively our results show joint effects of *EDM2*, *EDM3* and *IBM2* on basal immunity and overall plant growth to be genetically interlinked and functionally connected by their dependency on *SID2*, *PAD4* and *IBM1*. This view is further supported by effects we observed using the type III peroxidase inhibitor SHAM. The promotion of plant root and rosette growth by *EDM2*, *EDM3* and *IBM2* must be a consequence of their suppressive effects on basal immunity. Generally these three chromatin-associated regulators seem to contribute to a balanced defense-growth trade off by prioritizing overall plant growth over basal defense responses. Results of this study capture only a fraction of the roles *EDM2*, *EDM3* and *IBM2* have in coordinating immune mechanisms and developmental/growth-related processes. As outlined above, they also align the extent of basal defense with the timing of the floral transition and prioritize *RPP7*-mediated race-specific immunity over basal defense. Additional research is needed to uncover the mechanics of these coordinative processes and to understand the full scope and significance their have in mediating a balanced defense-growth trade off.

## Materials and Methods

### Plant Material and Growth Conditions

All the genotypes except specified are in Col-0 background. The following single mutants used in this study: *edm2-2* [4], *edm3-2* [17], *sg1-3/ibm2* [11], *sid2-2* [43], *pad4* [26], and *ibm1-4* [16]. The following double mutants used in this study: *edm2-2;edm3-2, edm2-2;sg1-3/ibm2, edm3-2;sg1-3/ibm2* were described previously [17]. The following transgenic lines used in this study: *EDM3S^np^-1*, *EDM3S^np^-2, EDM3L^np^-1*, *EDM3L^np^-2*, *IBM2S-1* and *IBM2L-1* were described previously [17], *NahG* [44]. The genomic *IBM1* region (*gIBM1*) or *gIBM1* with a deletion of the heterochromatic repeats in its 7th intron (*gIBM1ΔHR*) were cloned as described previously [9] and transformed into *edm2-2*, *edm3-2* and *sg1-3/ibm2*, respectively, to generate the *gIBM1* and *gIBM1ΔHR* transgenic lines. Double mutants were made by crossing and F2 seeds were screened by genotyping PCR to select homozygotes. Plants are grown either on ½ MS medium or in soil under long day (16h/8h light-dark cycles).

### Primary Root Length Measurement

Plants were grown vertically on ½ MS medium for five or seven days. Primary root length of the indicated genotypes was measured using ImageJ.

### Rosette Area Measurement

Seedlings were untreated or treated with 20μM SHAM (S607-5G, Sigma) at day 14 and day16. Rosette areas were pictured and measured using ImageJ at day 25.

### RNA Extraction and qRT-PCR

Total RNA of the aerial parts of two weeks old plants was extracted using Trizol reagent (Life technologies) according to the instructions of the manufacturer. 1μg of RNA was reversed transcribed into cDNA using Maxima Reverse Transcriptase (Fisher scientific). The cDNA products were used for Real-time PCR with CFX CONNECT detection system (Bio-Rad). All primers used are listed in Table 4.1.

### Peroxidase Activity Measurement and inhibitor experiments

Two weeks old seedlings were used for peroxidase activity measurement as described previously [45]. Effects of peroxidase inhibition were tested using salicylhydroxamic acid (SHAM),

### Hyaloperonospora arabidopsidis Infection Assay

*Hpa*Noco2 infection assays were performed as previously described [46].

## Acknowledgements

This work was supported by National Science Foundation (NSF) grant IOS-1457329 to TE.

## Supporting information captions

**S1 Fig.**
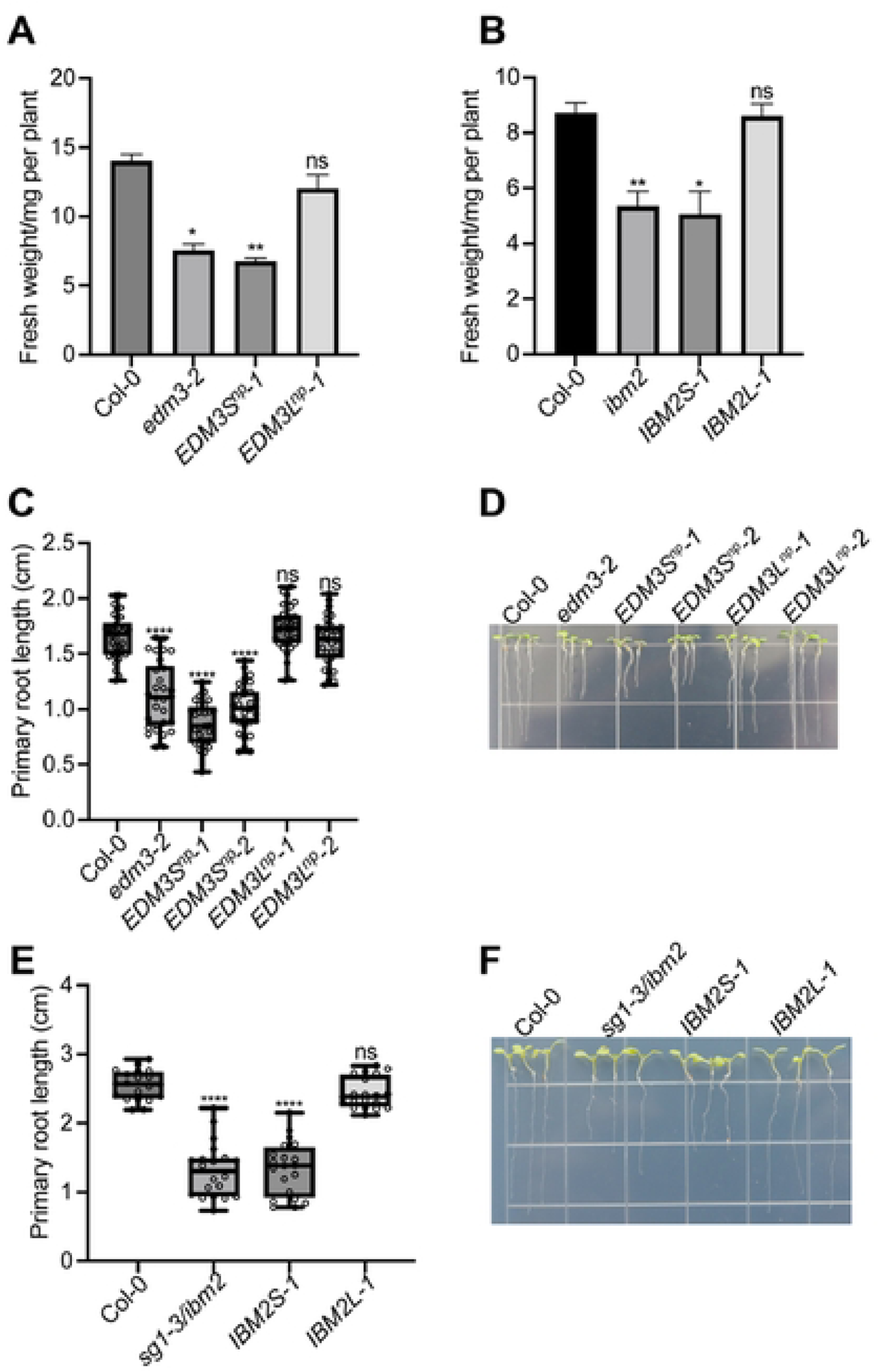
Only expression longer EDM3 and IBM2 isoforms can rescue growth defects in *edm3* and *ibm2* mutants. **A** Whole plan-fresh weight of 15-day-old Col-0, *edm3-2* or *EDM3* isoform-specific complementation lines (*EDM3S^np^-1*, *EDM3L^np^-1)* grown in soil. *EDM3S^np^-1* and *EDM3L^np^-1* express in the *edm3-2* mutant background either the short or long EDM3 isoform, respectively, driven by the native *EDM3* promoter (*np*). **B**. Fresh weight of 12-day old Col-0, *ibm2* and *IBM2* isoform-specific complementation lines (*IBMS-1*, *IBM2L-1)* grown in soil. *IBMS-1*, *IBM2L-1* express in the *ibm2* mutant background either the short or long IBM2 isoform, respectively, driven by the native *IBM2* promoter. The *sg1-3* mutant allele of *IBM2* was used for all experiments. **C & E.** Primary root length of 5-day-old plants of Col-0, *edm3-2* and *ibm2* plants as well as EDM3-and IBM2-isoform specific complementation lines grown on agar plates. **D & F.** Representative images of plants used in panels C and E. Data information: Error bars represent standard errors from three independent experiments. Asterisks indicate significant differences compared to Col-0 based on Student’s t-test. (*, p < 0.05; **, p < 0.01; ****, p < 0.0001; ns, no significance). n ≥ 28 (C) and n ≥ 20 (E).

**S2 Fig.**
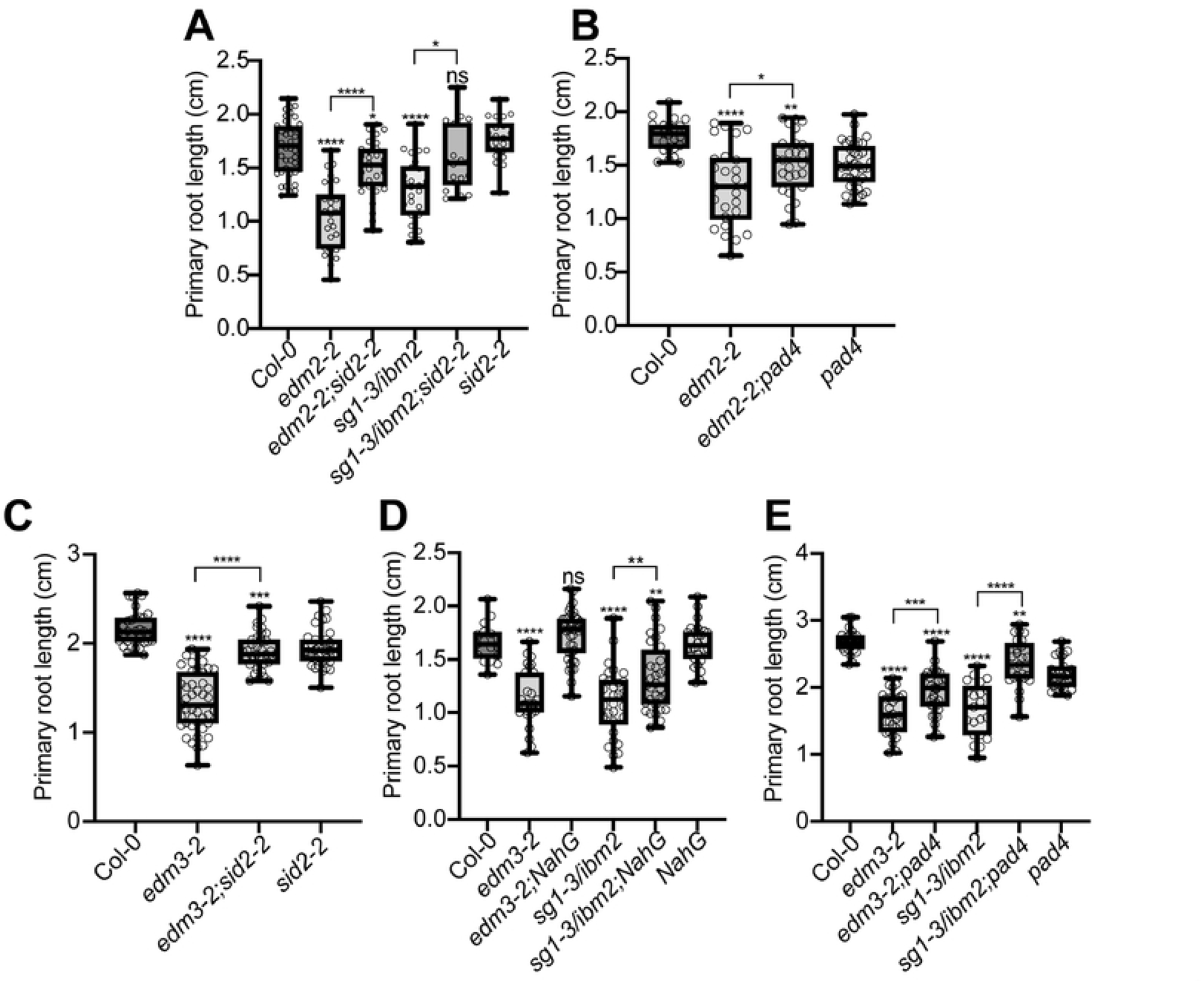
Primary root length of 5-day-old plants of the indicated genotypes. Primary root length of the indicated genotypes was measured using ImageJ. Data information: Data shown in each separate panel were generated simultaneously. Asterisks indicate significant differences compared to Col-0 based on one-way ANOVA (A-E). (*, p < 0.05; **, p < 0.01; ***, p < 0.001; ****, p < 0.0001; ns, no significance). n ≥ 18 (A), n ≥ 24 (B), n ≥ 33 (C), n ≥ 17 (D), n ≥ 21 (E).

**S3 Fig.**
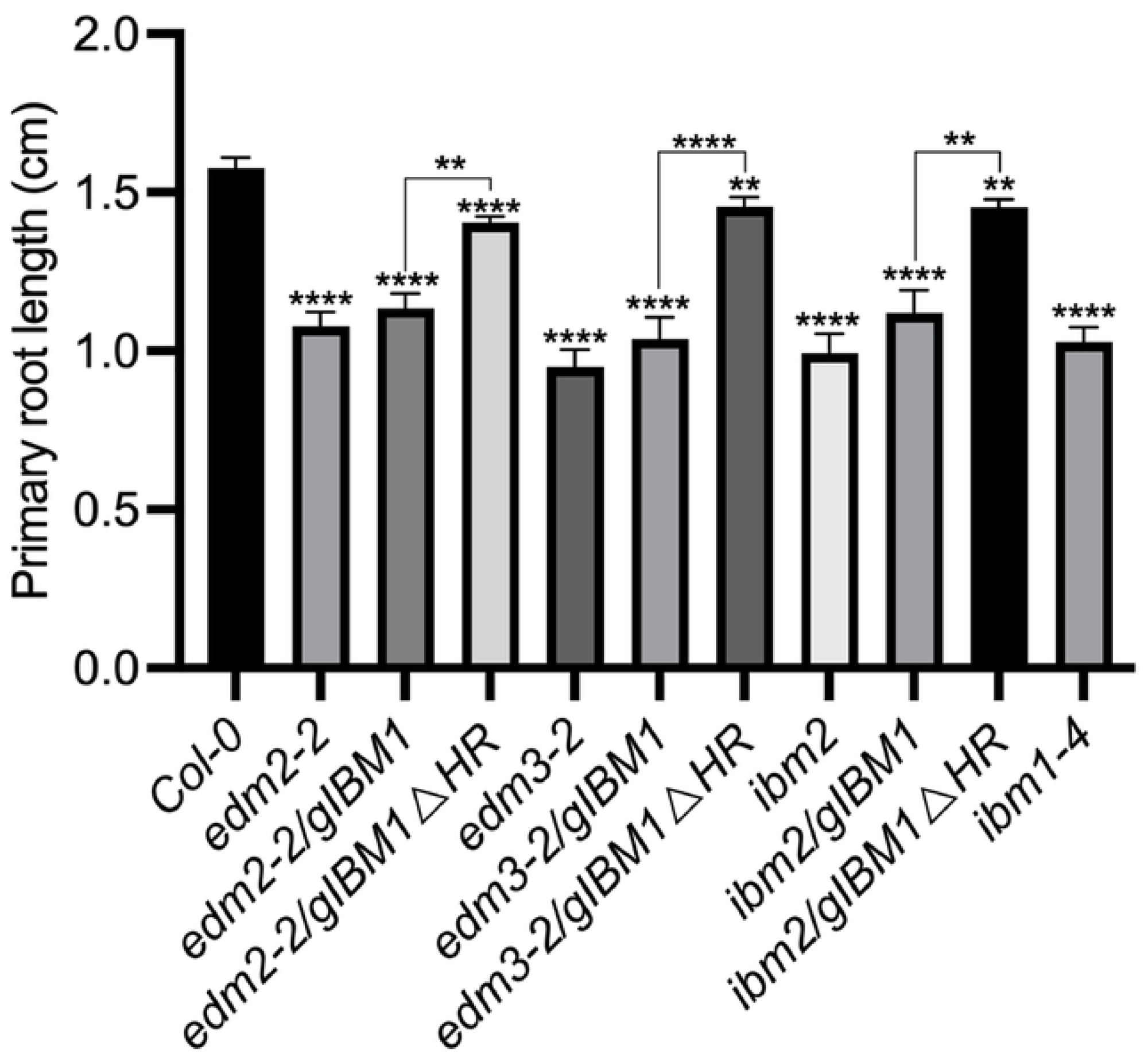
Short primary root length in mutants of *EDM2*, *EDM3* and *IBM2* was partly rescued by wild type expression of *IBM1L*. Primary root lengths of five day-old plants of the indicated genotypes were measured using ImageJ. Error bars represent standard errors. Asterisks indicate significant differences compared to Col-0 based on one-way ANOVA. (**, p < 0.01; ****, p < 0.0001). n ≥ 20.

**S1 Table. Primers used in this study**

## References

1. Karasov TL, Chae E, Herman JJ, Bergelson J. Mechanisms to Mitigate the Trade-Off between Growth and Defense. Plant Cell. 2017;29: 666–680.

2. Kempel A, Schädler M, Chrobock T, Fischer M, van Kleunen M. Tradeoffs associated with constitutive and induced plant resistance against herbivory. Proc Natl Acad Sci U S A. 2011;108: 5685–5689.

3. Züst T, Rasmann S, Agrawal AA. Growth–defense tradeoffs for two major anti-herbivore traits of the common milkweed Asclepias syriaca. Oikos. 2015. Available: https://onlinelibrary.wiley.com/doi/abs/10.1111/oik.02075

4. Eulgem T, Tsuchiya T, Wang X-J, Beasley B, Cuzick A, Tör M, et al. EDM2 is required for RPP7-dependent disease resistance in Arabidopsis and affects RPP7 transcript levels. Plant J. 2007;49: 829–839.

5. Tsuchiya T, Eulgem T. Co-option of EDM2 to distinct regulatory modules in Arabidopsis thaliana development. BMC Plant Biology. 2010. p. 203. doi:10.1186/1471-2229-10-203

6. Tsuchiya T, Eulgem T. The Arabidopsis defense component EDM2 affects the floral transition in an FLC-dependent manner. Plant J. 2010;62: 518–528.

7. Tsuchiya T, Eulgem T. Mutations in EDM2 selectively affect silencing states of transposons and induce plant developmental plasticity. Sci Rep. 2013;3: 1701.

8. Lei M, La H, Lu K, Wang P, Miki D, Ren Z, et al. Arabidopsis EDM2 promotes IBM1 distal polyadenylation and regulates genome DNA methylation patterns. Proc Natl Acad Sci U S A. 2014;111: 527–532.

9. Saze H, Kitayama J, Takashima K, Miura S, Harukawa Y, Ito T, et al. Mechanism for full-length RNA processing of Arabidopsis genes containing intragenic heterochromatin. Nat Commun. 2013;4: 2301.

10. Wang X, Duan C-G, Tang K, Wang B, Zhang H, Lei M, et al. RNA-binding protein regulates plant DNA methylation by controlling mRNA processing at the intronic heterochromatin-containing gene IBM1. Proc Natl Acad Sci U S A. 2013;110: 15467–15472.

11. Coustham V, Vlad D, Deremetz A, Gy I, Cubillos FA, Kerdaffrec E, et al. SHOOT GROWTH1 maintains Arabidopsis epigenomes by regulating IBM1. PLoS One. 2014;9: e84687.

12. Duan C-G, Wang X, Zhang L, Xiong X, Zhang Z, Tang K, et al. A protein complex regulates RNA processing of intronic heterochromatin-containing genes in Arabidopsis. Proc Natl Acad Sci U S A. 2017;114: E7377–E7384.

13. Tsuchiya T, Eulgem T. An alternative polyadenylation mechanism coopted to the Arabidopsis RPP7 gene through intronic retrotransposon domestication. Proc Natl Acad Sci U S A. 2013;110: E3535–43.

14. Deremetz A, Le Roux C, Idir Y, Brousse C, Agorio A, Gy I, et al. Antagonistic Actions of FPA and IBM2 Regulate Transcript Processing from Genes Containing Heterochromatin. Plant Physiol. 2019;180: 392–403.

15. Lai Y, Cuzick A, Lu XM, Wang J, Katiyar N, Tsuchiya T, et al. The Arabidopsis RRM domain protein EDM3 mediates race-specific disease resistance by controlling H3K9me2-dependent alternative polyadenylation of RPP7 immune receptor transcripts. Plant J. 2019;97: 646–660.

16. Lai Y, Lu XM, Daron J, Pan S, Wang J, Wang W, et al. The Arabidopsis PHD-finger protein EDM2 has multiple roles in balancing NLR immune receptor gene expression. PLoS Genet. 2020;16: e1008993.

17. Wang J, Eulgem T. The Arabidopsis RRM domain proteins EDM3 and IBM2 coordinate the floral transition and basal immune responses. Plant J. 2023. doi:10.1111/tpj.16364

18. Zhang Y-Z, Lin J, Ren Z, Chen C-X, Miki D, Xie S-S, et al. Genome-wide distribution and functions of the AAE complex in epigenetic regulation in Arabidopsis. J Integr Plant Biol. 2021;63: 707–722.

19. Bowling SA, Guo A, Cao H, Gordon AS, Klessig DF, Dong X. A mutation in Arabidopsis that leads to constitutive expression of systemic acquired resistance. Plant Cell. 1994;6: 1845–1857.

20. Dietrich RA, Delaney TP, Uknes SJ, Ward ER, Ryals JA, Dangl JL. Arabidopsis mutants simulating disease resistance response. Cell. 1994;77: 565–577.

21. Petersen M, Brodersen P, Naested H, Andreasson E, Lindhart U, Johansen B, et al. Arabidopsis map kinase 4 negatively regulates systemic acquired resistance. Cell. 2000;103: 1111–1120.

22. Clarke JD, Volko SM, Ledford H, Ausubel FM, Dong X. Roles of salicylic acid, jasmonic acid, and ethylene in cpr-induced resistance in arabidopsis. Plant Cell. 2000;12: 2175–2190.

23. Maleck K, Neuenschwander U, Cade RM, Dietrich RA, Dangl JL, Ryals JA. Isolation and characterization of broad-spectrum disease-resistant Arabidopsis mutants. Genetics. 2002;160: 1661–1671.

24. Wildermuth MC, Dewdney J, Wu G, Ausubel FM. Isochorismate synthase is required to synthesize salicylic acid for plant defence. Nature. 2001;414: 562–565.

25. Gaffney T, Friedrich L, Vernooij B, Negrotto D, Nye G, Uknes S, et al. Requirement of salicylic Acid for the induction of systemic acquired resistance. Science. 1993;261: 754–756.

26. Jirage D, Tootle TL, Reuber TL, Frost LN, Feys BJ, Parker JE, et al. Arabidopsis thaliana PAD4 encodes a lipase-like gene that is important for salicylic acid signaling. Proc Natl Acad Sci U S A. 1999;96: 13583–13588.

27. Parker JE, Holub EB, Frost LN, Falk A, Gunn ND, Daniels MJ. Characterization of eds1, a mutation in Arabidopsis suppressing resistance to Peronospora parasitica specified by several different RPP genes. Plant Cell. 1996;8: 2033–2046.

28. Wojtaszek P. Oxidative burst: an early plant response to pathogen infection. Biochem J. 1997;322 (Pt 3): 681–692.

29. Daudi A, Cheng Z, O’Brien JA, Mammarella N, Khan S, Ausubel FM, et al. The apoplastic oxidative burst peroxidase in Arabidopsis is a major component of pattern-triggered immunity. Plant Cell. 2012;24: 275–287.

30. Torres MA, Dangl JL, Jones JDG. Arabidopsis gp91phox homologues AtrbohD and AtrbohF are required for accumulation of reactive oxygen intermediates in the plant defense response. Proc Natl Acad Sci U S A. 2002;99: 517–522.

31. Bolwell GP, Wojtaszek P. Mechanisms for the generation of reactive oxygen species in plant defence – a broad perspective. Physiological and Molecular Plant Pathology. 1997. pp. 347–366. doi:10.1006/pmpp.1997.0129

32. Smirnoff N, Arnaud D. Hydrogen peroxide metabolism and functions in plants. New Phytol. 2019;221: 1197–1214.

33. Schopfer P. Hydrogen peroxide-mediated cell-wall stiffening in vitro in maize coleoptiles. Planta. 1996;199: 43–49.

34. Neuser J, Metzen CC, Dreyer BH, Feulner C, van Dongen JT, Schmidt RR, et al. HBI1 Mediates the Trade-off between Growth and Immunity through Its Impact on Apoplastic ROS Homeostasis. Cell Rep. 2019;28: 1670–1678.e3.

35. Greenberg JT, Ausubel FM. Arabidopsis mutants compromised for the control of cellular damage during pathogenesis and aging. Plant J. 1993;4: 327–341.

36. Yu IC, Parker J, Bent AF. Gene-for-gene disease resistance without the hypersensitive response in Arabidopsis dnd1 mutant. Proc Natl Acad Sci U S A. 1998;95: 7819–7824.

37. Frye CA, Tang D, Innes RW. Negative regulation of defense responses in plants by a conserved MAPKK kinase. Proc Natl Acad Sci U S A. 2001;98: 373–378.

38. Dangl JL, Dietrich RA, Richberg MH. Death Don’t Have No Mercy: Cell Death Programs in Plant-Microbe Interactions. Plant Cell. 1996;8: 1793–1807.

39. McDowell JM, Dangl JL. Signal transduction in the plant immune response. Trends Biochem Sci. 2000;25: 79–82.

40. Lorrain S, Vailleau F, Balagué C, Roby D. Lesion mimic mutants: keys for deciphering cell death and defense pathways in plants? Trends Plant Sci. 2003;8: 263–271.

41. Moeder W, Yoshioka K. Lesion mimic mutants: A classical, yet still fundamental approach to study programmed cell death. Plant Signal Behav. 2008;3: 764–767.

42. Lee Y, Rubio MC, Alassimone J, Geldner N. A mechanism for localized lignin deposition in the endodermis. Cell. 2013;153: 402–412.

43. Dewdney J, Reuber TL, Wildermuth MC, Devoto A, Cui J, Stutius LM, et al. Three unique mutants of Arabidopsis identify eds loci required for limiting growth of a biotrophic fungal pathogen. Plant J. 2000;24: 205–218.

44. Delaney TP, Uknes S, Vernooij B, Friedrich L, Weymann K, Negrotto D, et al. A central role of salicylic Acid in plant disease resistance. Science. 1994;266: 1247–1250.

45. Bindschedler LV, Dewdney J, Blee KA, Stone JM, Asai T, Plotnikov J, et al. Peroxidase-dependent apoplastic oxidative burst in Arabidopsis required for pathogen resistance. Plant J. 2006;47: 851– 863.

46. McDowell JM, Cuzick A, Can C, Beynon J, Dangl JL, Holub EB. Downy mildew (Peronospora parasitica) resistance genes in Arabidopsis vary in functional requirements for NDR1, EDS1, NPR1 and salicylic acid accumulation. Plant J. 2000;22: 523–529.

